# Theta activity supports landmark-based correction of naturalistic human path integration

**DOI:** 10.1101/2025.03.20.644367

**Authors:** Clément Naveilhan, Raphaël Zory, Klaus Gramann, Stephen Ramanoël

## Abstract

How do humans integrate landmarks to update their spatial position during active navigation task? Using immersive virtual reality and high-density mobile EEG, we investigated the neural underpinnings of landmark-based recalibration during path integration. Our findings reveal that a briefly presented intramaze landmark effectively corrected accumulated homing errors. However, this effect was transient and less optimal when participants were highly confident in their self-motion-based spatial representation suggesting that internal priors hinder the assimilation of novel spatial cues. At the neural level, RSC theta-band activity supported these recalibration processes. When fine adjustments of the spatial representation were sufficient, landmark presentation elicited stronger theta-band activity and greater phase resetting compared to when substantial spatial updating was necessary. Our results also revealed motor-related theta activity that scaled with acceleration during rotational corrections, highlighting the dual role of theta in the flexible integration of multimodal signals, involved in both landmark-based spatial updating and self-motion encoding.

## Introduction

Spatial navigation relies on the integration of multiple sources of idiothetic and allothetic information to enable efficient orientation and make it possible to reach the intended destination. When allothetic information such as visual cues are absent, individuals must rely exclusively on the continuous integration of idiothetic information stemming from self-motion including optic flow, vestibular, proprioceptive, and motor efference copy, to maintain a sense of position^1,2^. However, the reliability of self-motion cues diminishes over distance due to the gradual accumulation of sensory noise and leads to increased navigational errors, a limitation referred to as “leaky path integration“^3,4^. To recalibrate the path integrator and stop the accumulation of errors, visual information are used when available^5,6^. Such recalibration is central to the navigation process in humans, and visual landmarks play a pivotal role in anchoring spatial representations^7^. However, despite the pervasive nature of this intermodal recalibration, how the human brain integrates and combines external landmarks with self-motion cues remains elusive.

A substantial body of work has identified the brain regions involved in the isolated processing of either idiothetic or allothetic information. These studies have highlighted the role of the hippocampus in using self-motion cues for navigation, along with contributions from extrahippocampal regions^8–10^. Conversely, when landmarks are the only source of information during stationary spatial orientation tasks, studies reported the involvement of several high-level visual regions, including the parahippocampal place area (PPA), the retrosplenial cortex (RSC, also referred to as the Medial Place Area in the context of human scene processing^11,12^), and the occipital place area (OPA^13–15^). Although the distinct neural correlates associated with the integration of idiothetic and landmark information suggest separate mechanisms for navigation based on self-motion and visual cues, these processes are instead considered as complementary^16^. Self-motion cues provide fundamental spatial information regarding position and orientation, whereas external visual cues recalibrate the path integration system, reducing errors and enhancing overall navigational accuracy^17–21^. In this context, recent studies using animal models highlighted the key role of the RSC region for combining these two sources of information for efficient spatial navigation^22–26^. The integration of information from self-motion and landmark-based cues is facilitated by the connectivity of the RSC with cortical and subcortical regions like the hippocampus, anterior thalamic nuclei, parahippocampal structures, and the visual system^27^. This hub role played by the RSC supports the binding of allothetic and idiothetic cues into flexible spatial representations that accommodate shifts in sensory input and context-dependent navigation demands^28–30^.

In human navigation, previous studies have proposed that the role of the RSC region is mainly related to visuospatial processing of landmarks^31,32^. It has also been extended to the positional coding of self-motion cues as a way to keep track of translation and rotation^33–36^. However, these studies have predominantly used stationary experimental protocols and have not considered several self-motion cues (*i.e.,* vestibular information, proprioception, motor efference copies) that are crucial in naturalistic human navigation^37–39^. Advances in EEG and virtual reality technologies now make it possible to study brain dynamics during unrestricted movement in controlled environments^40–42^. Recent investigations using these methods have expanded the role of the RSC region to active human spatial navigation. They highlighted for example its involvement in the computation of full-body rotations, as reflected in slow-frequency oscillations, such as theta activity^36,43–45^. Despite these initial findings, it remains unclear how the human brain dynamically combines the information conveyed by self-motion cues and landmarks to update the path integration system in naturalistic human navigation.

To address this issue, we combined virtual reality and high-density mobile EEG to study how participants perform a path integration task relying solely on self-motion cues. Along the path, participants periodically reported the remembered starting position and their confidence level in their response by pointing to the start location and subsequently indicating their confidence on a virtual scale. Once per path, after their pointing response, they were briefly presented with a previously seen landmark to help correct their estimate of the starting location. Consistent with prior findings^4^, we observed a progressive accumulation of errors throughout the path integration process. The presentation of the landmark allowed a transient improvement in accuracy, but this corrective effect was reduced when participants exhibited high confidence in their prior spatial estimates. Theta-band activity in the RSC reflected the recalibration process. When the landmark aligned with predictions from self-motion cues, increased theta activity and phase resetting provided optimal conditions to facilitate the integration of both cues supporting fine adjustment of the representation. During the physical rotation to correct the homing, theta power also increased. However, rather than scaling with the magnitude of correction, it was modulated by participant acceleration. These findings underscore a dual role for low-frequency oscillations in the RSC, mediating both landmark-based recalibration of spatial representation and the encoding of self-motion cues during naturalistic human navigation.

## Results

Data were collected from 28 young adults performing a path integration task in a virtual reality environment combined with high-density mobile EEG (128 electrodes). Each path included a sequence of four stops, after which participants returned to the starting location (**Figure 1.A**). At each stop along the path, they physically turned to indicate their starting position and rated their confidence in their response. Then, a previously seen proximal landmark was presented at either the second or third stop of the sequence. Participants had to wait without moving for 1 second after presentation of the landmark and then update their prior estimates based on the newly available visual information (**Figure 1.B**; see **Methods**). To disentangle the distinct contributions of translations and rotations to error accumulation during path integration, we systematically manipulated these parameters across three different paths configurations (**Figure 1.E,** see **Methods** for more details).

**Figure 1.**
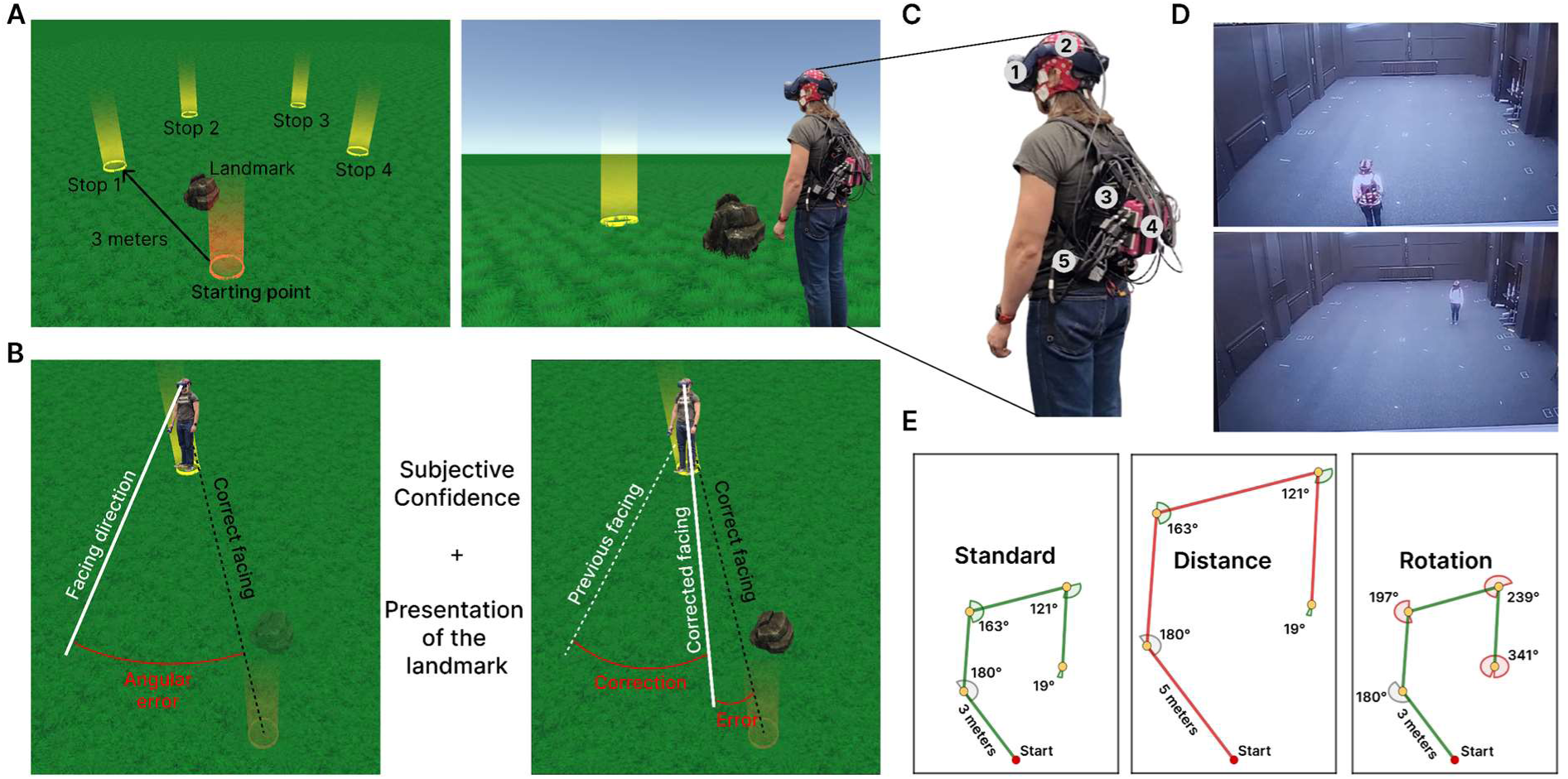
Experimental protocol and setup. **A.** Top-down view of the environment *(left)* and the first-person perspective experienced by participants *(right)*. **B.** Sequence of the actions performed at a stop during landmark presentation. Participants pointed towards their starting position *(left)*, then the landmark appeared for one second while they remained still *(middle)* and then rotated to correct their pointing *(right)*. Only absolute errors were analyzed. **C.** Experimental setup worn by participants, including (1) HTC Vive Pro virtual reality headset, (2) 128-channel ANT Neuro EEG cap, (3) Zotac computer, (4) two cascading ANT Neuro amplifiers, and (5) two lithium batteries. **D.** The 8 × 12 m dark room, showing a participant at the start of a trial (*top*) and at the final stop (*bottom*) before returning to the starting point. **E.** The three path configurations used in the experiment to selectively manipulate the amount of translation and rotation.

Participants were equipped with a fully mobile virtual reality and EEG setup (**Figure 1.C**), enabling them to move freely within an 8 × 12-meter room devoid of any sensory cues that could serve as landmarks (**Figure 1.D**). We spatially filtered the EEG data and subsequently source reconstructed the origin of the recorded brain dynamics focusing our analyses on the RSC and including data from 23 participants (**Figure 5**).

### Landmarks allow for resetting the path integration system

In this initial behavioral analysis, we extracted participants’ pointing errors defined as the deviation between their pointing response to the starting location and the correct homing angle (hereafter referred to as “pointing error” for simplicity). We analyzed this error across the four different stops and three path configurations to assess how the landmark reduced the pointing error and whether this recalibration effect persisted over subsequent path segments (**Figure 2.A**). After comparing models using the Akaike Information Criterion (AIC)^46^, we applied a linear mixed model with the following parameters :

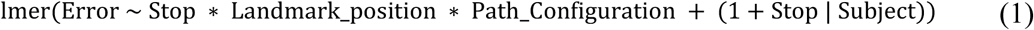

**Figure 2.**
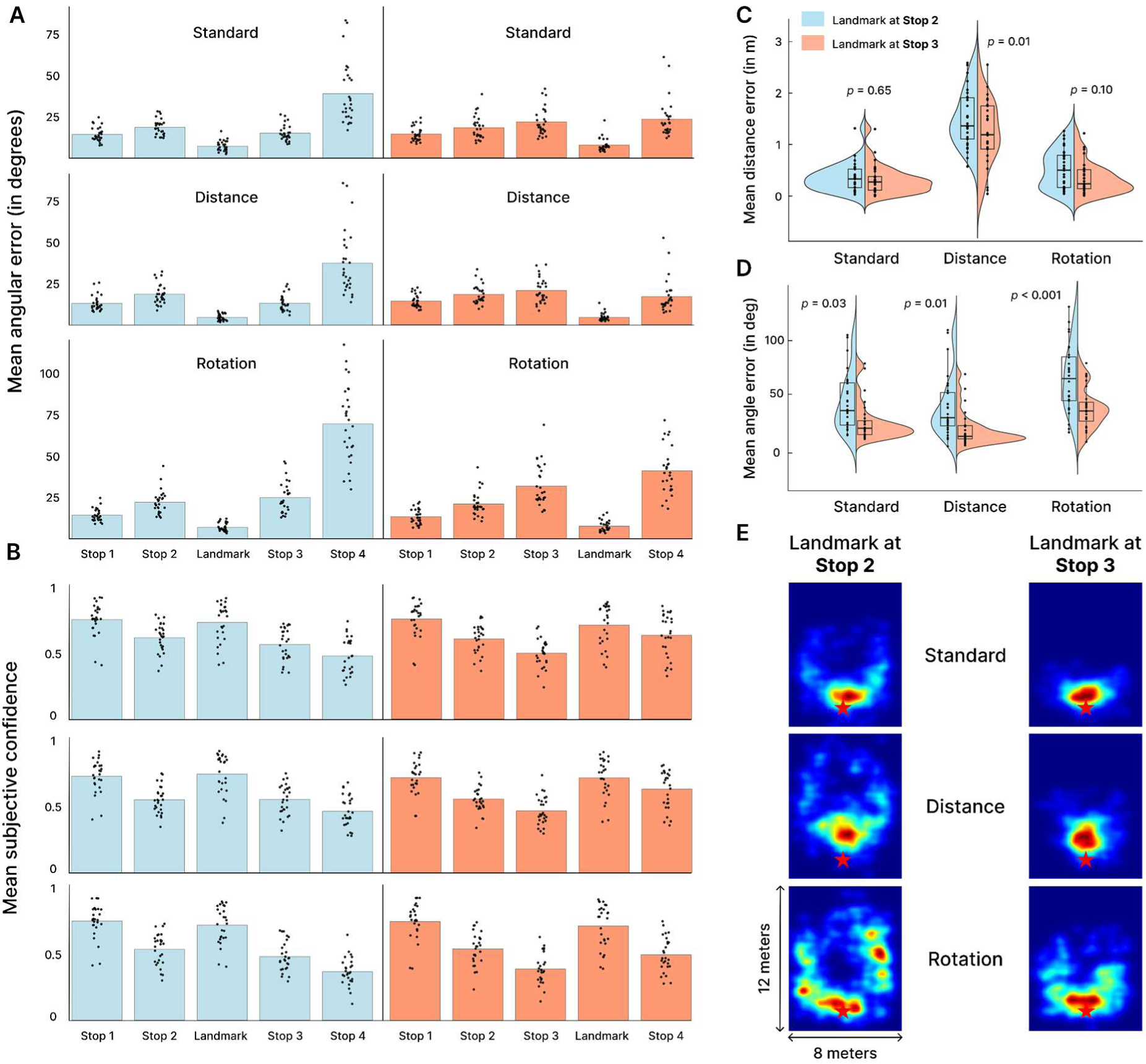
Behavioral results. **A.** Homing pointing error at each stop in the sequence and after andmark presentation. Light blue represents landmarks shown at the second stop, and coral represents those shown at the third stop. Lines indicate different path configurations (standard, distance, and rotation). **B.** Participant confidence ratings on a visual scale from 0 (not confident) to 1 (extremely confident). **C.** Absolute distance error for the homing segment. **D.** Mean pointing error for the fourth stop. **E.** Density plot of homing positions averaged across all participants and experimental conditions. The red star marks the starting position participants were instructed to return to.

In this model, we analyzed the pointing *Error* including as fixed factors: *Stop*, representing the stop at which participants performed the pointing task (1st, 2nd, 3rd, or 4th); *Landmark_position*, indicating the stop where the landmark was presented (2nd or 3rd stop); and *Path_configuration*, reflecting the navigation condition (standard, long-distance, or increased-rotation). Additionally, *Stop* was included as a random slope and intercept for each subject.

We found a significant main effect of stop position in the path sequence (F_(3,38.55)_ = 24.51, *p* < 0.001, η_p_^2^ = 0.66, 95% CI = [0.45, 0.76]), with pointing error increasing from the first stop (Estimated Marginal Means : M = 14.9; SD = 1.08) to the second stop (M = 20.4; SD = 1.38; (t_(28)_ = 4.46, *p* < 0.001, d = 0.57, 95% CI = [0.31, 0.84]). This finding replicates Stangl *et al.* (2020)^4^ results and supports a leaky path integrator with a rapid accumulation of error in human navigation in the absence of visual landmarks to anchor the spatial representation. Notably, participants already demonstrated varying pointing errors at the first stop, despite walking in a straight line and simply needing to turn and point backward 180°. Since this condition did not require them to recall the starting position but only to execute a physical rotation, this suggests that a significant portion of the pointing error arises from the rotational movement itself, sometimes referred to as execution error^47–49^. For the first two stops, no effect of landmark presentation should be expected since the landmark was not presented until stop 2 or 3. The results confirmed the absence of an impact of landmark presentation (all *p* > 0.71) or path configuration (all *p* > 0.25), indicating consistency across pre-landmark conditions. However, at the third stop, pointing errors were significantly reduced in the case where a landmark had been presented at the previous stop (t_(560)_ = 4.41, *p* < 0.001, d = 0.68, 95% CI = [0.36, 0.98]), compared with pointing errors when no landmark had been presented, suggesting that the landmark correction persisted to some extent at the subsequent stop.

Next, we examined pointing errors at the fourth stop and found that they were significantly larger when the landmark was presented at the second stop (M = 53.1 ± 4.16) than when the landmark was presented at the third stop (M = 30.7 ± 4.16; t_(560)_ = 15.31, *p* < 0.001, d = 2.32, 95% CI = [1.99, 2.66]). This suggests that while landmarks can temporarily reduce accumulated noise, the noise rapidly builds up again, indicating that the corrective effect seems to be short-lived.

To confirm that the landmark effectively reduced the accumulated error and to examine how the path configuration and the stop at which the landmark was presented (either Stop 2 or 3) influenced this correction, we applied the following model:

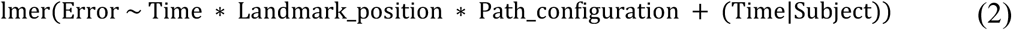

In this model, the factor *Time* corresponds to the moment when the pointing error was calculated (*i.e.,* before or after landmark presentation). Our results confirmed that the landmark effectively reduced pointing errors (F_(1,28)_ = 156.09, *p* < 0.001, η_p_^2^ = 0.85, 95% CI = [0.73, 0.90]), with pointing errors decreasing from 23.49 ± 1.45 degrees before presentation to 7.46 ± 0.84 degrees after. This correction effect differed depending on whether the landmark appeared at the second or third stop (F_(1,280)_ = 13.09, *p* < 0.001, η_p_^2^ = 0.05, 95% CI = [0.01, 0.10]), driven by larger pre-landmark pointing errors in the third stop condition (t_(280)_ = 5.13, *p* < 0.001, d = 0.78, 95% CI = [0.47, 1.09]). However, pointing errors after the presentation of the landmark did not differ between stop conditions (t_(280)_ = 0.03, *p* = 0.974). These findings suggest that regardless of when the landmark was presented or the path configuration (all *p* > 0.058), participants successfully calibrated their position and orientation in space based on the landmark.

### RSC Theta activity correlates with the extent of landmark-based correction

To explore the neural basis of the observed landmark-based recalibration of the path integration system, we analyzed how theta activity in the RSC regions was modulated by the presentation of a landmark (**Figure 5**). Theta band activity was functionally defined between 2 and 8 Hz, following previous research on human navigation^50–53^. Activity was examined over the period during which the participants were presented with the landmark and had to remain still for 1 second before rotating to adjust their pointing response.

Analysis of z-scored theta activity in the RSC revealed two distinct peaks (**Figure 3.A 3.B**). The first peak occurred around 300 ms after the landmark was presented, while the second peak appeared at the onset of participants’ rotational movements to correct their previous pointing, following the 1-second stationary period. Notably, theta activity in the RSC for the first peak exhibited a linear decrease as a function of correction magnitude (mean beta within the significant window, 264–412 ms: –0.193, *p*_FDR_ < 0.05; **Figure 3.C**). In our model, we included path configuration as a co-variate to control for its potential influence on the analysis. However, no significant effect was observed at any time point after false discovery rate (FDR) correction (all *p*_FDR_ > 0.84), indicating that path configuration did not affect theta activity after landmark presentation. At this point, it is important to clarify how we interpret the magnitude of correction and how it reflects the degree of alignment between landmark information and self-motion cues. Small corrections suggest that the spatial representation was already well-aligned with self-motion cues, requiring only minor adjustments of this representation before anchoring for later retrieval. In contrast, larger corrections indicate that the landmark prompted a substantial readjustment of the spatial representation. In the former case, smooth recalibration to stable values remains possible, a mechanism proposed by computational models to be supported by phase reseting^54^, underscoring the critical role of theta phase in spatial coding (*e.g.,* the oscillatory interference model).

**Figure 3.**
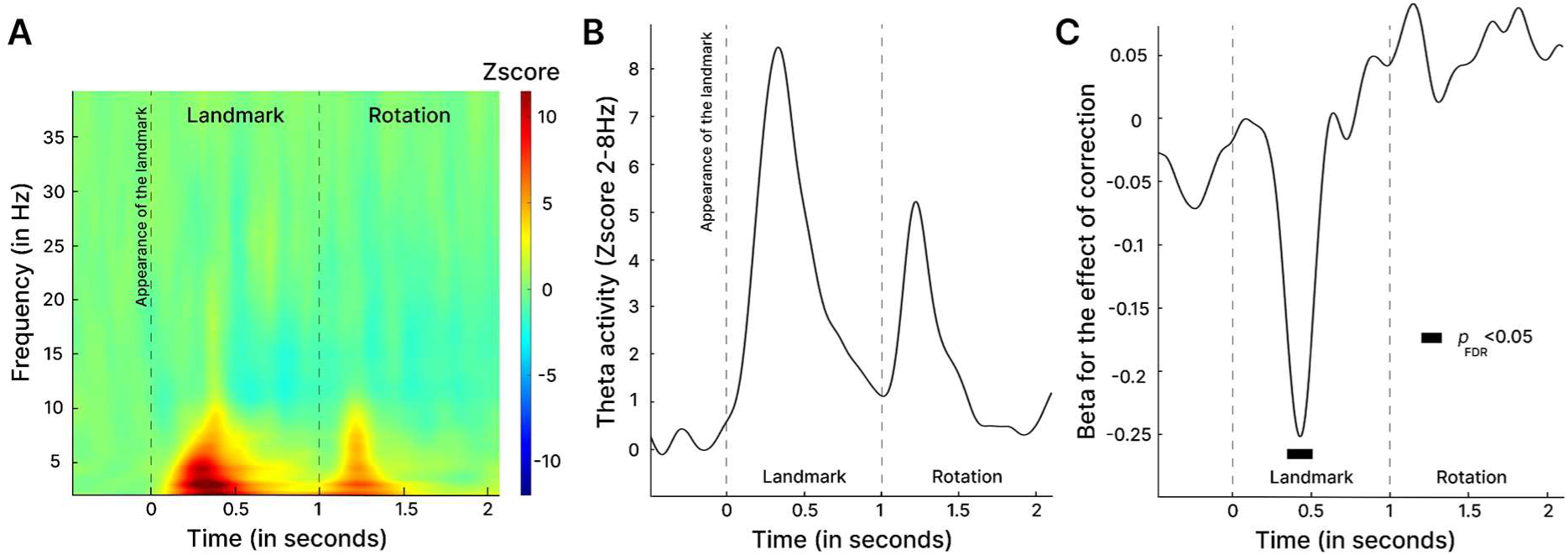
EEG results for the power analysis. **A.** Z-scored time-frequency representation (2-40Hz) aligned to the onset of landmark appearance and the physical rotation required for correction. **B.** Mean Z-scored theta activity (2–8 Hz) within the same time window. **C.** Beta estimates of the effect of correction on theta activity. Black regions indicate statistically significant clusters identified through permutation testing with FDR correction.

To investigate this possible relationship between theta activity and landmark-based path integration updating, we analyzed intertrial phase coherence (ITPC)^55^ in the RSC. To this end, we separated and compared trials based on the median correction value (**Figure 4**) to ensure an equal number of trials, thereby controlling for potential biases in the estimates^56^. Results of the cluster-based permutation tests indicated that, following landmark presentation, ITPC values were higher when participants made smaller corrections compared to larger corrections (*p*_cluster_ < 0.05, for the 94-550 time window). This effect was specific to theta activity, with a significant cluster observed between 90 and 554 ms after landmark presentation, while no difference was found during the rotation period. These results suggest that when the landmark information aligned with the prediction of participants based on self-motion cues (*i.e.,* their pointing before landmark presentation)—RSC theta activity exhibited greater power and phase coherence. In contrast, when participants required to update their spatial representation, theta power and phase coherence were reduced, a result that we can interpret as a decrease in phase resetting^57–59^. This phase alignment effect, observed when the landmark matched participants’ predicted spatial representation, was confirmed by the polar histogram of theta phase (**Figure 4.E**), with an increased concentration of phases between 200° and 270°. Statistical analyses of the mean resultant vector length (MRL; **Figure 4.F**) indicated greater phase angle consistency when the landmark elicited small corrections but only in the time window after landmark presentation (F_(1,23.62)_ = 14.36, *p* <0.001, η_p_^2^ = 0.38, 95% CI = [0.09, 0.60]; with no effect in the two other time windows, all *p* > 0.145). The oscillatory interference model^60^ suggests that phase resetting plays a crucial role in correcting path integration errors^61^. According to this model, environmental cues such as landmarks, turning points, and boundaries provide a resetting signal that synchronizes all oscillators to the same phase. Our findings offer evidence for this mechanism in naturalistic human navigation. Specifically, they propose that phase resetting increases when allothetic and idiothetic information align, allowing fine adjustments of the spatial representation before being anchored for later recall.

**Figure 4.**
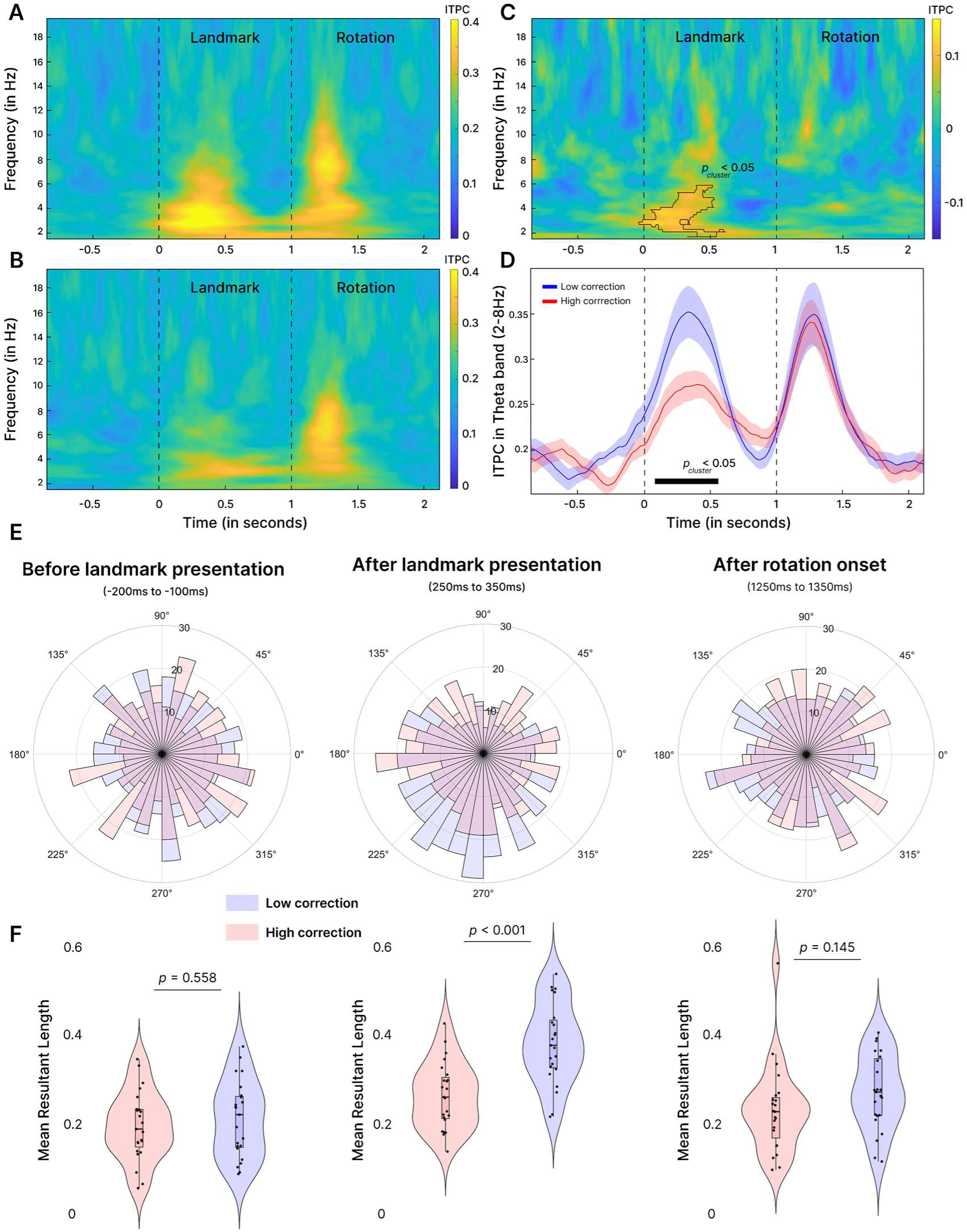
EEG results for the phase analyses. **A.** Intertrial phase coherence for trials in which the landmark elicited a small correction, indicating that the landmark information was aligned with the path integration-derived representation **B.** Same as A., but for trials when there was an important misalignment between each representations. **C.** Difference between high- and low- correction trials. The black outline indicates cluster-based significant differences. **D.** Theta band results (2–8 Hz) with standard error. The black bar indicates significant clusters from permutation testing between the two conditions, corrected for false discovery rate (FDR). **E.** Polar histogram of theta phase distribution relative to correction magnitude, presented across three 100 ms time windows: before and after landmark presentation, and at a matched interval after rotation onset. **F.** Violin plot illustrating the mean resultant length, an index of phase angle consistency. Statistical comparisons for each time window were performed using a linear mixed model.

Our results also revealed a second peak of theta activity occurring when participants initiated the rotation. However, this activity was not related to the magnitude of correction, suggesting that theta activity during physical rotation may originate from a motoric component. This interpretation aligns with the proposed multifaceted role of low-frequency oscillations in spatial cognition, supporting their involvement in both spatial memory processes and the integration of body-related sensory information during movement^62^. Building on this framework and recent studies in rodents reporting an association between acceleration and theta activity^63^, we also extracted both peak speed and peak acceleration from participants within a 500 ms time window following the onset of rotation, along with the corresponding theta activity in the RSC. We then applied linear mixed-effects models to examine the relationships between theta activity and these kinematic parameters. The results indicated that theta activity increased linearly with peak acceleration (*β* = 0.563, 95% CI = [0.139, 0.988], t_(848)_ = 2.604, *p* = 0.009). In contrast, no such relationship was observed for peak speed (t_(848)_ = 0.31, *p* = 0.752). **Supplementary 1** provides extended results with control analyses supporting the movement-related origin of the second peak of RSC theta activity reported here.

### The degree of confidence modulates the integration of landmark-based information

We aimed to determine whether participants’ confidence prior to landmark presentation influenced how they adjusted their pointing (**Figure 2.B**). We first assessed their ability to accurately judge their own performance using the visual scale, computing confidence-weighted accuracy (CWA which ranges from 0 to 1; see Methods for details). Results showed that participants effectively used subjective confidence measures and accurately judged their performance, although this ability declined over successive stops (see **Supplementary 2** for detailed analyses).

We then investigated how the degree of confidence before the landmark presentation influenced the extent to which participants integrated landmark information to update their spatial representation. Specifically, we hypothesized that when participants are highly confident in their spatial representation based on self-motion cues, they would be less likely to update their response in light of added information available. Based on model comparisons using the AIC, we selected a linear mixed model with the following parameters:

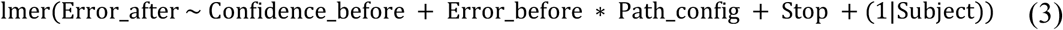

Similarly to previous analysis, we found that pointing errors after landmark presentation remained consistently low across the different stops in the sequence (*p* = 0.411), confirming that participants successfully used the landmark across the different stops. However, these pointing errors after landmark presentation were significantly modulated by the pointing errors before landmark presentation (*β* = 0.068, 95% CI [0.042, 0.094], *p* < 0.0001), indicating that although participants adjusted their pointing upon landmark presentation, their pointing estimates were still influenced by their prior estimates. In line with our hypothesis, we observed more pronounced pointing errors in post-landmark pointing as a function of pre-landmark confidence (*β* = 1.158, 95% CI [0.132, 2.181], *p* = 0.027), suggesting that higher confidence prior to landmark presentation was associated with greater residual error afterward, independently of the stop (*p* = 0.307) or the configuration of the path (*p* = 0.078). This finding implies that when participants reported a higher initial confidence they relied less optimally on landmark information for correction, a result that we discuss in the following discussion in the context of Bayesian optimal cue weighting^64,65^.

### Presentation of landmarks improves performance of outbound path, but only for long distances

Although our primary focus in this study was on homing pointing errors, participants also completed a homing path after the fourth stop by walking back to the remembered starting location. From this, we calculated the distance homing error^66^, which is the error in total distance independent of any angular error :

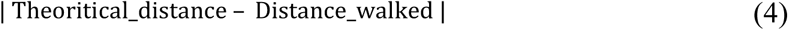

Our results (**Figure 2.C**) indicate that path configuration significantly influenced distance error (F_(2,140)_ = 107.07, *p* < 0.001, η_p_^2^ = 0.60, 95% CI = [0.51, 0.68]), with greater distance homing errors observed for paths with longer legs (M = 1.36 ± 0.06) compared to both the normal (M = 0.34 ± 0.06) and long-rotation (M = 0.44 ± 0.06) conditions. However, no significant difference was found between the long-rotation and normal path conditions (t_(140)_ = 1.31, *p* = 0.39), where travel distances were matched. These results indicate that longer walking distances during path integration increase final distance errors, while larger rotation angles, as previously reported, primarily drive rotational errors. This pattern suggests that adding noise to either the rotational or translational components of path integration produces corresponding errors, supporting the existence of distinct systems for tracking translation and rotation^34,67^. Additionally, the location of landmark presentations significantly influenced distance homing errors (F_(1,140)_ = 7.28, *p* = 0.008, η_p_^2^ = 0.05, 95% CI = [0.00, 0.14]), with larger errors observed when the landmark was presented at the second stop (M = 0.80 ± 0.04) compared to the third stop (M = 0.63 ± 0.04). However, this effect was specific to the long-leg condition (t_(140)_ = 2.56, *p* = 0.011, d = 0.67, 95% CI = [0.15, 1.20]), with no significant differences observed in the standard or increased rotation conditions (*p* > 0.1). These results suggest that the proximal landmark also provided distance-related information, which enhanced performance specifically when participants had to traverse longer distances. The signed distance error values indicated a systematic undershooting of the starting point in the long-leg condition, with participants exhibiting an average error of –1.33 ± 0.55 meters, compared to 0.05 ± 0.40 meters in the standard condition (t_(140)_ = 1.39, *p* < 0.001, d = 3.69, 95% CI = [3.15, 4.22]), and 0.16 ± 0.43 meters in the increased rotation condition (t_(140)_ = 1.51, *p* < 0.001, d = 3.99, 95% CI = [3.49, 4.50]).

## Discussion

In this study, we investigated the neural dynamics underlying path integration and visual landmark integration during naturalistic human spatial navigation. Our results suggest that while an intramaze landmark effectively reduces path integration error, angular homing errors rapidly accumulate thereafter. Using high-density mobile EEG (128 electrodes) combined with spatial filtering and source reconstruction, we identified theta activity in the RSC as a key marker of this recalibration process. When only fine adjustments were required to correct the idiothetic spatial representation using landmark information, we observed increased theta power and phase resetting. In contrast, when landmark triggered the need for a more extensive remodeling of the spatial representation we observed a less pronounced phase resetting. Although participants successfully used the landmark, this correction effect was modulated by their confidence in the spatial representation derived from idiothetic cues. Specifically, higher confidence prior to the presentation of the landmark was associated with suboptimal integration of new information, suggesting that stronger priors reduce reliance on incoming sensory cues. Our findings present convincing evidence for a dual function of theta activity, highlighting a distinct motor-related theta response at the initiation of rotation, triggered by peak acceleration. This dual role underscores the importance of theta activity in the RSC for the dynamic integration of multimodal signals, contributing to both landmark-based spatial updating and vestibular self-motion encoding.

Consistent with our initial hypothesis, we found that regardless of the path configuration or the stop position in the sequence, participants successfully corrected their homing responses following the landmark presentation. However, angular homing errors rapidly re-accumulated at the next stop and during the outbound path (*i.e.,* returning to the starting position after the fourth stop), resembling the accumulation observed before the landmark was presented^4^. Using a path integration task with four stops, we were able to directly track both error accumulation and landmark-based correction for the first time. The transient nature of the landmark correction suggests that the majority of accumulated error arises from noise and velocity gain bias, while memory decay of the starting position plays only a minimal role^4^. During the outbound path, participants exhibited greater pointing errors in conditions with increased rotations and greater distance homing errors when required to walk farther. These findings suggest that selectively increasing noise in either the rotational or translational component of path integration led to corresponding errors, supporting the idea of distinct systems that track translation and rotation. This interpretation aligns with the organization of the vestibular system^68^ and is further supported by evidence that rotational errors increase specifically in the early stages of Alzheimer’s disease, whereas distance errors appear to be more related to a general process of aging^69,70^. Crucially, landmark presentation reduced both types of error, indicating that the landmarks provided not only directional but also positional information—a property typically attributed to intramaze landmarks, in contrast to distal landmarks, which primarily convey stable directional cues^71–73^. Interestingly, the landmark selectively corrected the specific type of error introduced in each experimental condition. For example, when distance was increased, participants performed the return path more accurately when the landmark was presented closer in time to the homing response (*i.e.,* at the third compared with the second stop), whereas no improvement in accuracy was observed for homing pointing errors. These findings further support the idea that translation and rotation are distinct components of path integration, with the RSC proposed to track both but prioritizing the most task-relevant information^34,67^. Finally, analysis of the signed distance error revealed that participants systematically underestimated long distances, a finding also consistent with previous research^5,74,75^. This result may be partially explained by a regression to the mean, as participants in the long-distance condition tended to produce distances closer to those required in the two other conditions (*i.e.,* standard and rotation) which were thus presented more often^5,76^.

Regarding neural correlates of the path integration correction upon landmark presentation we reported a burst of theta activity around 300ms after landmark presentation in a cluster of independent components with their equivalent dipole models located in or near the RSC. This finding extends previous results from the rodent literature to human participants regarding the role of the RSC for processing the most stable features in the environment, which, under natural conditions, are typically landmarks^24,27,31,77^. Human electrophysiology results are more sparse, but the role of slow frequency oscillations during landmark processing as a means of spatial updating has been proposed based on intracranial EEG recorded from the hippocampus (HPC) as a means of spatial updating^78,79^. Using this invasive approach in rodents, hippocampal theta oscillations were reported to be phase-locked to activity in the dysgranular subregion of the RSC^80^, highlighting the critical role of RSC-HPC connectivity in spatial cognition^81,82^. Crucially, we reported not only a burst of theta activity upon landmark presentation, but also a scaling of this activity dependent on the required correction of the homing representation. This activation pattern was further coupled with an increase in theta phase alignment when the landmark information aligned with the subjective spatial representation based on self-motion cues. This concurrent increase in theta power (**Figure 3**), inter-trial phase coherence and phase alignment (**Figure 4**) can be interpreted as a phase reset of oscillations in the RSC^57–59^. In other words, when landmark information align to some extent the spatial representation formed through self-motion cues, RSC theta activity exhibit increased phase resetting. Theta phase has been proposed to play a key role in memory and spatial navigation^83,84^, with phase resetting interpreted as a mechanism to support optimal conditions for synaptic potentiation at the neural level^85,86^. In our context, this mechanism may serve to facilitate the correction of cumulative error through environmental sensory input^60,87^, like proposed by attractor models, before anchoring the correct current spatial representation for later recall^84,88,89^. Supporting this interpretation, Güth *et al.* (2024)^90^ demonstrated that theta phase resetting is crucial for encoding salient spatial information during navigation. Our results extend these findings by reporting a correlation with the amount of spatial representation correction elicited by this information, confirming fMRI findings suggesting that RSC may contain spatial representation for both landmarks and self-motion cues^91^.

From a more mechanistic perspective, our findings extend the framework proposed by Campbell *et al.* (2018)^17^ in mice to humans. Here we propose that theta activity in the RSC may reflect the model proposed with the sub-critical and super-critical regimes of attractor network dynamics during spatial navigation. In the sub-critical regime, when discrepancies between path integration and landmark representations are minimal, we observed increased theta-band power and phase resetting compared to when the correction was larger. This enhanced neural synchronization could reflect the continuous, fine adjustments maintained by the attractor network to align internal spatial representations with external cues, supporting stable and accurate navigation as predicted by Campbell *et al.* (2018)^17^. Computational modeling by Monaco *et al.* (2011)^54^ has already suggested that a single cue can effectively reset oscillator phases, enabling corrections for both systematic errors and continuous noise in path integration^61^. However, this phase alignment may only be possible when the mismatch between both cues is minimal, as reported in our results with small corrections, and in previous results with a smooth resetting of the head direction system to stable values^92^. In cases of larger discrepancies, also referred to as the super-critical regime^17^, these fine corrections though phase resetting may not be sufficient. Thus, the appearance of the landmark triggers the need for significant internal reworking of the spatial representation, with a remodeling of the head direction system^92^ and a possible shift in the hippocampal representation^93^ before the corrected information is transferred across different brain regions through theta activity^94–97^. In this case of a large required correction we observed lower theta activity and phase alignment in the RSC, a finding that may reflect the delay associated with the necessary reprocessing of spatial representations. We speculate that this remapping may be associated with increased connectivity between the RSC and the hippocampus, but intracranial recordings would be necessary to explore this possibility in greater depth. Taken together, our non-invasive neural results provide compelling evidence for the role of theta activity in the RSC of freely moving humans in the landmark-based updating of spatial representations initially derived from path integration. These findings align with recent mechanistic models and further support the proposal that the RSC is involved in generating a prediction (in this case, based on self-motion cues and corresponding to the initial pointing response) and subsequently updating this prediction using newly available information, such as a landmark^22^.

Beyond landmark processing, we also observed increased theta activity in the RSC during participants’ physical rotation when they adjusted their angular homing pointing. However, this second theta peak was not correlated with the magnitude of the correction, suggesting that it is unrelated to the landmark updating process. Instead, we reported a linear scaling of activity in this frequency band with the peak acceleration in the 500 ms time window following participant onset of rotation time but no association with speed. These results replicate previous rodent findings during translational movement^63^ and extend these findings to mobile noninvasive recordings in humans. This also aligns with previous mobile EEG and fMRI studies proposing that the RSC plays a key role in computing heading rotation information, likely reflecting its involvement in the processing of head direction signals. As a central hub in the head direction cell network, the RSC is thought to support the integration of self-motion cues for maintaining a stable sense of orientation during navigation^34,36,44,45,67,98^. Thus our results strongly support that low-frequency oscillations have a multifaceted role^99^, with theta activity related to both the processing of landmark information and the encoding of vestibular information related to rotational acceleration.

Finally, after each homing pointing response along the paths, participants reported their subjective confidence. We found that they effectively used the confidence scale to evaluate their performance^100^, although this ability decreased along the different stops in the path. This result suggests that participants can judge their performance reasonably well, but that this judgment is still impacted by noise accumulation. Subjective confidence has been proposed to reflect a Bayesian representation of probability in the cortex^101,102^. The Bayesian framework provides a principled approach for optimally integrating multiple sensory cues, particularly under conditions of noisy or ambiguous sensory input^103–107^. By combining prior knowledge with sensory likelihoods to compute posterior distributions, Bayesian models enable optimal perception, resolve ambiguity, and generate reliable estimates for spatial navigation despite variable inputs^64,65,108–112^. Within this framework, it has been suggested that greater confidence in prior knowledge reduces the influence of new, discrepant information^113^. Consistent with this idea, we found that when participants were more confident in their pointing prior to landmark presentation they relied less on landmark information to adjust their response, leading to less effective corrections. Our results provide evidence in spatial navigation of how prior confidence influences cue weighting in a Bayesian framework, paving the way for further investigations that take advantage of this easy-to-use metric of subject confidence. Specifically, our results demonstrate that greater confidence in prior estimates reduces the weight assigned to new sensory cues, resulting in suboptimal adjustments, even when the landmark provides highly reliable information as in our protocol.

To conclude, our findings provide among the first non-invasive neuroimaging evidence that theta oscillations support landmark-based correction of path integration during naturalistic human navigation. While participants used landmarks to recalibrate spatial representation, this correction was modulated by prior confidence, highlighting the influence of internal priors on sensory integration. Theta phase resetting in the RSC facilitated spatial representation correction when fine adjustments of the spatial representation were sufficient. These findings provide compelling evidence supporting emerging mechanistic models of spatial updating. We also observed a second distinct theta peak during rotation which was driven by acceleration rather than homing correction, highlighting the multifaceted role of theta activity in human spatial navigation. Overall, our results underscore the power of mobile EEG in capturing the neural mechanisms underpinning dynamic spatial cognition, offering new insights into how the human brain flexibly integrates multimodal signals to support navigation in naturalistic settings.

## Methods

### Participants

We recorded the EEG activity of 29 healthy young participants with no history of neurological pathology and normal or corrected-to-normal vision. Participants signed an informed consent form approved by the Ethics Committee of the Technical University of Berlin (BPN_GRA_231204, Institute of Psychology & Ergonomics, Technical University of Berlin, Germany). They were recruited locally via an online platform and received monetary compensation (€12/hour) for their participation. We excluded 1 participant from our sample due to excessive artifacts in the EEG recording, resulting in a final sample of 28 participants (11 females, mean age: 25.92 ± 3.75). Participants completed a motion sickness questionnaire to ensure they reported no symptoms^114^, and a computerized version of the Perspective Taking Task / Spatial Orientation Test to assess their visuospatial abilities in a 2D environment^115^.

### Stimuli and procedure

The virtual environment was created using the Unity3D game engine (version 2021.3.8f1 for Windows) from Unity Technologies, San Francisco, California, USA. Rendering was performed using an HTC Vive Pro head-mounted display with a 90 Hz refresh rate, dual 3.5-inch AMOLED screens with 1440×1600 pixel resolution, 615 ppi, and a 110° nominal field of view. The HTC Vive Pro was connected to a VR-ready backpack computer (Zotac PC) equipped with an Intel 7th Gen Kaby Lake processor, GeForce GTX 1060 graphics, 32GB DDR4-2400 memory support, and running Windows 10 OS (ZOTAC Technology Limited, Fo Tan, Hong Kong). This setup was battery powered and remotely controlled. The participant’s head movements were recorded using an integrated HTC Lighthouse motion tracking system, which included four cameras operating at a sampling rate of 90 Hz and covering an area of 8 × 12 m.

The virtual environment consisted of an infinite floor with a grass texture and some added grass tufts to allow the integration of optic flow by the participants (**Figure 1.A**). The task consisted of an extended triangle completion task with 4 stops, each marked by a yellow ring that appeared along the path. At each stop, participants were asked to turn in the direction indicated by an arrow (either in the shortest direction in the normal and long-leg condition or in the longest in the long-rotation condition) in order to face their (unmarked) starting point as accurately as possible. They then had to press a button on the controller to validate their answer and were presented with a bar. They had to indicate their subjective confidence (*i.e.,* the longer the bar, the more confident they were). They were carefully familiarized with the subjective confidence indicator beforehand to ensure that they understood it well. Participants had to walk from one ring to another without any visual cues. After reaching each ring, they were presented with an arrow for 1 second, which indicated the direction they had to rotate. At either the second or third stop, we presented them with a landmark (*i.e.,* a stone) that they had seen at the beginning of the trial. Participants had to wait one second to look at the landmark before being asked to report again the position of their starting point and their subjective confidence, using the spatial information conveyed by the landmark to correct their previous pointing (**Figure 1.B**). After the 4 stops, participants then had to physically return to their starting point, validate their response, and report their confidence in their homing task. Finally, they were presented with a red ring indicating the true position of the starting point and had to walk to this ring to start a new trial.

The task included three different conditions, with trial order counterbalanced so that half of the trials began on the left and half on the right side of the physical space. We used three path configurations: a standard path with legs of three meters each, a long-leg path in which the length of each leg was increased to five meters, and an increased-rotation condition in which participants had to perform rotations at a larger angle (**Figure 1.C**). In addition, we included two landmark configurations in which the landmark was presented at either the second or the third stop. This design resulted in a total of 12 unique paths derived from the combination of 2 start positions (left/right) x 2 landmark positions (second stop/third stop) x 3 path types (standard/long leg/increased rotation). Each of these 12 unique paths was presented in a randomized order within each run, resulting in a total of 60 paths per participant, divided into 5 runs presented in a pseudorandomized order, each consisting of 12 paths to complete. On average, each run lasted 14 minutes, resulting in a total average recording time of 71.32 minutes. To maximize the informativeness of the landmark we ensured its stability (the position remained constant across trials), its usefulness for navigation (the position was close to the starting point), and its visual features (it conveyed radial information and was easily visible at different stops)^31^. We used a proximal landmark, similar to that used by Doeller *et al.* (2008)^116^ to ensure that the landmark information dominated over idiothetic information during path integration, as speculated in previous work^111^. This also allowed us to disregard a complex part of cue combination, namely the process that allows us to check the stability of a landmark before relying on the conveyed information^31,77,117^.

### EEG acquisition and processing

EEG signals were acquired at a sampling rate of 500 Hz using two cascaded eego™ mylab amplifiers (ANT neuro bv, Hengelo, Netherland) and digitized at 24-bit resolution (**Figure 1.D**). Wet electrodes were embedded in a 128-channel actively shielded Waveguard™ cap according to the extended 5% system^118^. The online reference electrode was CPz, and the ground electrode was a wired gel droplet. The impedances of all electrodes were carefully kept below 10 kΩ using conductive gel, with most electrodes below 5 kΩ. One electrode was placed below the participant’s left/right eye to improve the identification of eye-movement related activity in the subsequent processing steps. EEG recordings, stimulus presentation, and motion capture were synchronized using Lab Streaming Layer (LSL) software^119^ and recorded via the LabRecorder.

EEG data were preprocessed offline (**Figure 5**) using Matlab (R2024a; The MathWorks Inc., Natick, MA, USA) and the BeMobil pipeline^120^ working with scripts for the EEGLAB toolbox version 2024.0^121^. First, non-experimental time segments were removed, and we manually inspected the raw signal to remove excessive artefacts. The data were then downsampled to 250Hz, before line noise was removed using Zapline Plus^122^ which enables the automatic detection and removal of artifactual peaks (mainly 50Hz and 90Hz corresponding to line noise and the refresh rate of the head-mounted display). Bad electrodes were rejected using *clean_raw_data* with a correlation criterion of 0.8 and a maximum time of broken electrodes of 0.5. These electrodes were then interpolated using spherical spline interpolation, and the data were re-referenced to a common average reference. Subsequently, we cleaned the data in the time domain using ASR with a burst criterion of 30^123^, which resulted in the rejection of an average of 3.01 ± 3.76 percent of the data points. The data were then temporarily high-pass filtered at 1.75 Hz before AMICA decomposition^124^ was applied to decompose the mobile EEG data into statistically maximally independent components (ICs). Data points that did not fit the ICA model were rejected in 10 iterations using the AMICA algorithm with a sigma threshold of 3^125^. Finally, we fitted equivalent dipole models to each resultant IC using DipFit before labeling the identified components using the lite version of Iclabel^126^. We rejected components labeled as muscle, line noise, eye, and heart, and kept only those classified as at least 30% brain and with a residual variance (RV) of the respective ICs below 15%^127^. This resulted in a conservative average retention of 16.69 ± 6.89 components, in line with what has been reported in other studies using similar mobile EEG^43,45^. The data were then filtered between 0.3 and 50 Hz for further analysis, and epoched from 3 s before the landmark appeared to 4 s after. Finally, epochs containing artifacts greater than 100 μV were removed.

**Figure 5.**
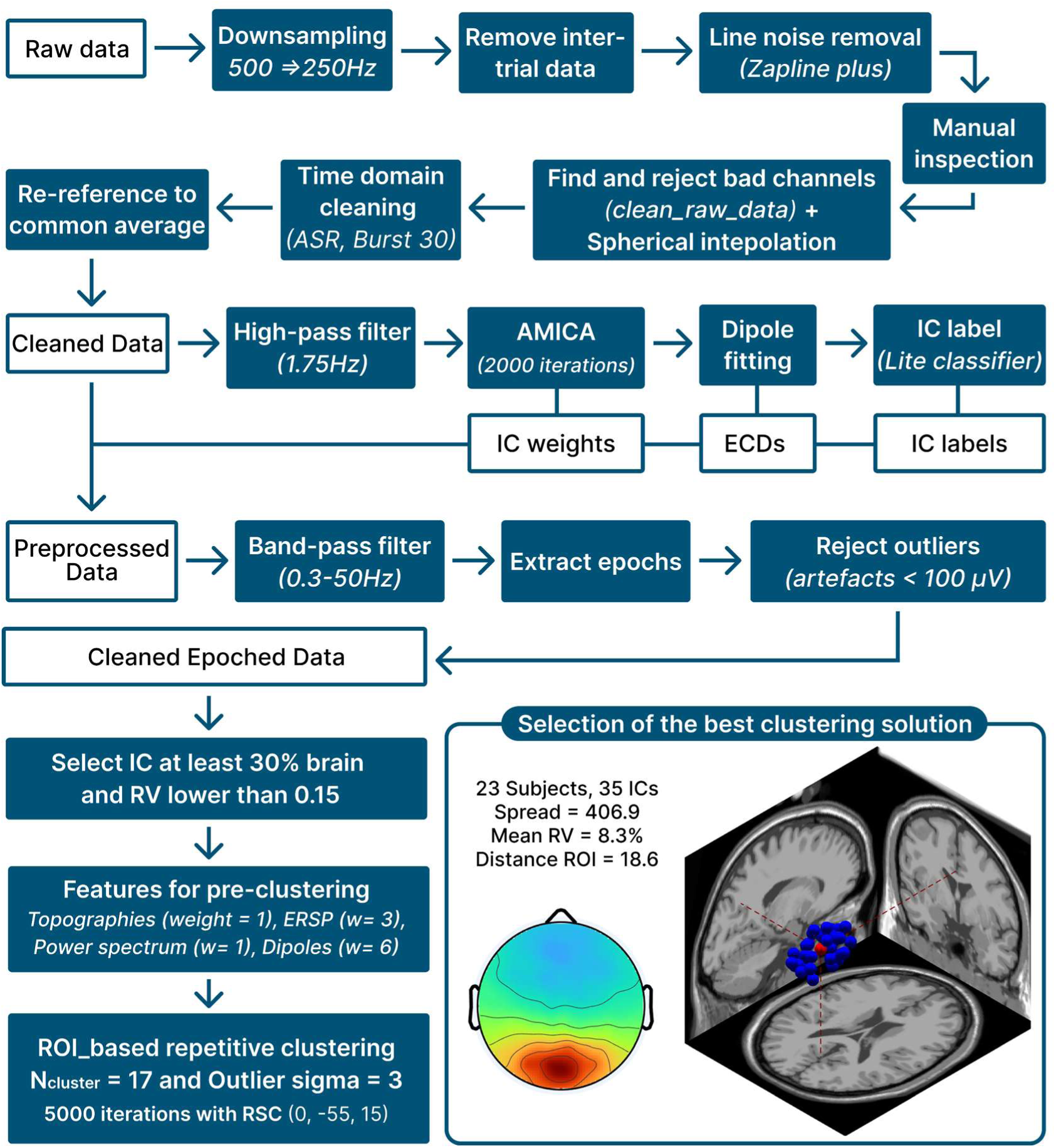
Presentation of the pipeline used for EEG preprocessing. EEG preprocessing and clustering pipeline. Raw data were downsampled (500 → 250 Hz), line noise removed (Zapline plus), and bad channels interpolated (clean_raw_data, spherical interpolation). Time-domain cleaning (ASR, burst = 30) was applied, followed by re-referencing and high-pass filtering (1.75 Hz). Independent component analysis (AMICA, 2000 iterations) was performed, with dipole fitting and component labeling. Preprocessed data were band-pass filtered (0.3–50 Hz), epoched, and outliers removed (<100 µV). ICs with ≥30% brain contribution and residual variance (RV) <0.15 were clustered using features (topographies, ERSP, power spectra, dipoles) and ROI-based repetitive clustering (N =17, 5000 iterations). The best clustering solution was selected based on spatial spread, mean RV, and ROI distance.

We then extracted the retained ICs and used repeated measures clustering with a region of interest (ROI) constraint. To do this, we first performed a pre-clustering principal component analysis using event-related spectral perturbations (ERSP, weight = 3), scalp topographies (weight = 1), the power spectrum (weight = 1), and the location of the dipole (weight = 6). We then performed 5000 iterations of k-mean clustering with 17 clusters and an outlier threshold of 3 standard deviations. We defined a region of interest centered around the retrosplenial complex using [0, −55, 15] as previous work suggested these coordinates to be the most representative^43^. Each clustering solution was then scored using a weighted combination of 6 metrics: the number of subjects per cluster (weight = 6), the number of IC per subject in the same cluster (weight = −3), the normalization of the spread (weight = −1), the mean RV (weight = −1), the distance from the ROI (weight = −3), and the Mahalanobis distance from the median of the multivariate distribution (weight = 1). We then combined and scaled these multivariate matrices on a scale from 0 to 1 and finally selected the highest ranked solution. If multiple ICs per subject were present in the cluster, we selected the best IC for each subject (based on dipole position and residual variance). Finally, we performed a time-frequency decomposition of the activity of the selected ICs in the cluster. To ensure the detection of transient activity, we used the Superlet approach^128^ implemented in the Fieldtrip toolbox^129^, and performed a single-trial z-score normalization as suggested by Grandchamp & Delorme^130^.

#### Behavioral and EEG analyses

Statistical analyses of the behavioral data were performed using the R statistical software (version 4.4.2, R Foundation for Statistical Computing, Vienna, Austria) with R studio (version 2024.09.1) and linear mixed effects models from the lme4 package^131^. To compare the different models, we computed the Aikaike Information Criterion (AIC)^46^ and selected the model with the lowest AIC value. To extract *p-*values, we used the *anova* function and the Satterthwaite method with type III sum of squares. Estimated marginal means (EMMs, referred to as M in the manuscript) were calculated using the *emmeans* package in R and are the mean values reported hereafter. Finally, post-hoc Tukey’s honestly significant difference (HSD) tests were performed, and partial eta-squared and Cohen’s d were reported as post-hoc effect size measures. To check for normality of residuals and homoscedasticity, both were carefully examined using quantile-quantile plots and box plots, respectively. For metacognitive abilities, we computed a composite score of normalized subjective confidence and normalized error:

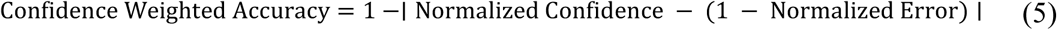

For EEG analysis, we used a hierarchical linear modeling approach^132^, with a linear mixed model implemented in custom Matlab scripts for the first level analysis. We then used 10 000 permutations to shuffle the error scores across participants and applied an FDR correction to the extracted *p*-values to examine the effects of the correction on theta activity. For intertrial phase coherence (ITPC), we separated epochs for each subject based on the median of their correction values. We then computed the Fourier spectrum using wavelets (width = 6; Gaussian width = 3) between 2 and 20 Hz with a 0.2 Hz step, and calculated the ITPC using the formula:

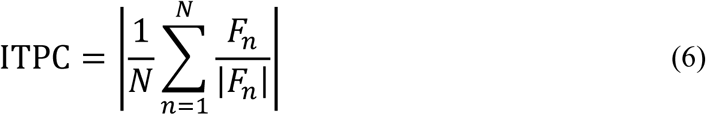

where N is the number of trials and F is the Fourier coefficient for the n^th^ trial. We then performed cluster-based permutation testing^133^ as implemented in the Fieldtrip toolbox, with 10 000 permutations.

For the mean resultant vector length (MRL) we used the following formula :

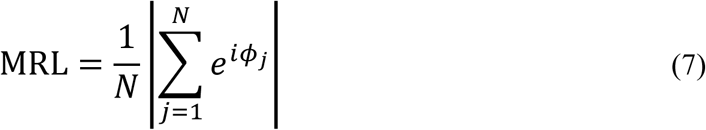

where N represents the number of trials and *e*^i*ϕ*j^ denotes the complex representation of the phase angle *ϕ*_j_ for each trial, as defined by Euler’s formula: *e*^i*ϕ*^ = cos(*ϕ*) + *i* sin(*ϕ*). Thus, the MRL value is a resultant vector, whose length reflects phase consistency: a longer resultant vector indicates stronger phase-locking, while a shorter vector signifies greater phase variability.

## Author Contributions

**Clément Naveilhan**: Conceptualization; Formal analysis; Investigation; Methodology; Writing—Original draft; Writing—Review & editing. **Raphaël Zory:** Funding acquisition; Writing—Review & editing. **Klaus Gramann**: Conceptualization; Methodology; Project administration; Writing—Review & editing. **Stephen Ramanoël**: Conceptualization; Funding Acquisition; Methodology; Project administration; Supervision, Writing—Review & editing.

## Acknowledgements

This research would not have been possible without the generous help of our volunteer participants, and the authors are truly grateful for their support. We also thank Catherine Buchanan for her careful reading of the manuscript and her feedback.

This work was supported by the French government through the France 2030 investment plan managed by the National Research Agency (ANR), as part of the Initiative of Excellence Université Côte d’Azur under reference number ANR-15-IDEX-01 and, in particular, by the interdisciplinary Institute for Modeling in Neuroscience and Cognition (NeuroMod) of Université Côte d’Azur.

## Data and code availability

All the raw data and the analysis codes generated for the present study are available online on the OSF repository of the study.

## Supplementary materials

### Supplementary 1

In this analysis, we investigated how the kinematic parameters of participants’ rotations influenced RSC theta activity. Rotational data along the Y-axis were preprocessed to compute the first and second derivatives, corresponding to speed and acceleration, within a 500-ms window following the onset of rotation. We extracted the peak and mean values of speed and acceleration in this window, along with the corresponding peak and mean theta activity in the RSC. As reported in the Results section, theta activity was significantly associated with peak acceleration (*β* = 0.563, 95% CI = [0.139, 0.988], t_(848)_ = 2.604, *p* = 0.009). This effect was further supported by permutation testing, in which theta activity values were shuffled 10 000 times across participants (N_permutation_ = 10000, *p* = 0.010). In contrast, no significant relationship was observed for mean values of either speed (*p* = 0.262) or acceleration (*p* = 0.398). The absence of an effect for mean values may reflect the transient, peaked nature of rotational acceleration, which is not captured by averaging over the 500 ms window. To ensure the robustness of the peak-based analysis, we compared these results, where values were extracted from a 3-point window centered on the peak, to values extracted using 1-point and 5-point windows. We found no effect of window size on the relationship between acceleration and theta activity (F_(2,2544)_ = 0.001, *p* = 0.999), supporting the generalizability of our approach.

Finally, we confirmed that the acceleration-theta relationship was absent during a 500-ms window following landmark presentation (*p* = 0.644). This result is consistent with participants remaining still as instructed while viewing the landmark, as indicated by the near-zero rotational speed (0.051 ± 0.06 rad/s equivalent to 2.92 ± 3.44 deg/s) during this period.

### Supplementary 2

In this analysis, we assessed whether participants accurately judged their pointing performance and examined whether the accuracy of these judgments was influenced by specific parameters manipulated in our protocol:

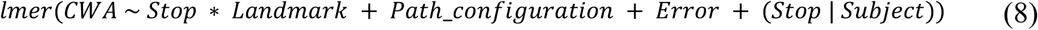

After excluding three participants who did not use the scale correctly (one consistently reported a value of 1, and two consistently reported a value of 0), the estimated marginal mean of the intercept was 0.698 (SE = 0.0181, 95% CI [0.661, 0.735], t_(26)_ = 38.580, *p* < 0.0001), indicating that, on average, participants were able to accurately assess their pointing performance. Permutation testing further confirmed that their metacognitive judgments were significantly better than chance (mean permutation = 0.614 ± 0.003, *p* < 0.001, N_permutation_ = 10,000). Interestingly, participants’ metacognitive accuracy was modulated by their position within the stop sequence (F_(3,41.67)_ = 21.66, *p* < 0.001, η_p_^2^ = 0.61, 95% CI = [0.40, 0.73]). Specifically, they judged their performance more accurately at the first stop (M = 0.817 ± 0.03) than at the second (M = 0.690 ± 0.02; t_(26.3)_ = 6.50, *p* < 0.001, d = 1.61, 95% CI = [1.09, 2.13]), third (M = 0.620 ± 0.02; t_(26.4)_ = 7.90, *p* < 0.001, d = 2.50, 95% CI = [1.83, 3.16]), or fourth stop (M = 0.666 ± 0.02; t_(32.2)_ = 6.88, *p* < 0.001, d = 1.92, 95% CI = [1.34, 2.51]). At the fourth stop, metacognitive accuracy was higher when the landmark was presented at the third stop (M = 0.709 ± 0.02) compared with landmark presentation at the second stop (M = 0.623 ± 0.02; t_(562)_ = 6.18, *p* < 0.001, d = 1.09, 95% CI = [0.72, 1.45]). Thus, these results suggest that as participants progressed along the path, they integrated increasingly noisy and ambiguous sensory information, leading to an increase error accompanied by a reduced ability to accurately judge their performance.

